# Monitoring lysosomal catabolism: a sensitive probe for assessing targeted lysosomal degradation of extracellular proteins

**DOI:** 10.1101/2024.10.18.619124

**Authors:** Ryan W. Holly, Yi-Zhen Hu, Vaishnavi Beena Valsan, Christos Kougentakis, Nektaria Petronikolou, Tao Chen, Diana Li, Aimei Chen, Steve Rank, Kyle H. Doran, Eric Turtle, Christopher Langsdorf, Matthew J. Shurtleff

## Abstract

Extracellular targeted protein degradation (eTPD) is an emerging therapeutic field. The Lysosome targeting chimera (LYTAC) is a therapeutic modality that promotes degradation of extracellular drivers of disease in the lysosome. While widely available pH-sensitive probes may report on lysosome delivery, these probes do not necessarily report on the enzymatically active functional state of the lysosome. We report the development and application of a sensitive fluorescent probe, LysoLight Deep Red, to monitor catabolism of internalized proteins in the lysosome based on cleavage by cathepsin proteases. We demonstrate the application of Lysolight Deep Red to monitor the catabolic fate of therapeutic monoclonal antibodies, ASGPR-targeted LYTAC therapeutics and LYTAC targets in immortalized cell lines and in primary human hepatocytes.

## Introduction

Contemporary antibody-based therapeutics employ a wide variety of formats and mechanisms of action (MOA) beyond a simple target binding mechanism. These approaches include antibody-dependent cellular cytotoxicity (ADCC), cis or trans co-targeting using bispecific antibodies, targeted toxin delivery using antibody drug conjugates (ADC), radioimmunoconjugates and extracellular targeted protein degradation (eTPD) utilizing a proximity inducing MOA (Reviewed in: [1]). Extracellular targeted protein degradation (eTPD) is a recently described therapeutic approach that utilizes cellular machinery to degrade circulating or membrane proteins that drive disease (Reviewed in: [2,3]). As an example, LYTACs are proximity-inducing heterobifunctional molecules that consist of 1) a binder to a target of interest, 2) a binder to an internalizing receptor (*e.g.*, CI-M6PR, ASGPR) and 3) a linker connecting the two binding moieties[4,5]. Highlighting the interest in eTPD as a therapeutic approach, a wealth of additional modalities have been described that operate through the same basic principle of induced proximity between a target and a cell surface protein resulting in lysosomal degradation of the target (e.g. Sweeping antibodies, Seldegs, IFLDs, DENTACs, MoDE-As, KineTACs, AbTACs, ROTACs)[6–13].

Both ADC toxin delivery and eTPD approaches require internalization of the drug and delivery of the drug and/or target to the lysosome. However, binding of an antibody to a cell surface protein does not invariably lead to antibody internalization or catabolism. Antibodies and their associated targets can potentially enter a recycling pathway rather than a degradation pathway which may compete with the desired MOA. Therefore, when developing biologic-based therapies it may be important to evaluate the cellular localization and ultimately catabolic fate of the therapeutic and target. While fluorescent tools exist to monitor internalization of extracellular proteins into the endocytic pathways (e.g. pH-sensitive dyes)[14], measuring the catabolic fate of internalized molecules remains time consuming, low throughput and semi-quantitative (e.g. immunoblotting). Here, we report the development and application of a small molecule dye that can be conjugated to any lysine-containing protein and remains non-fluorescent until the probe reaches the lysosome and is cleaved by lysosomal cathepsins resulting in a bright fluorescent signal (Figure 1A).

**Figure 1:**
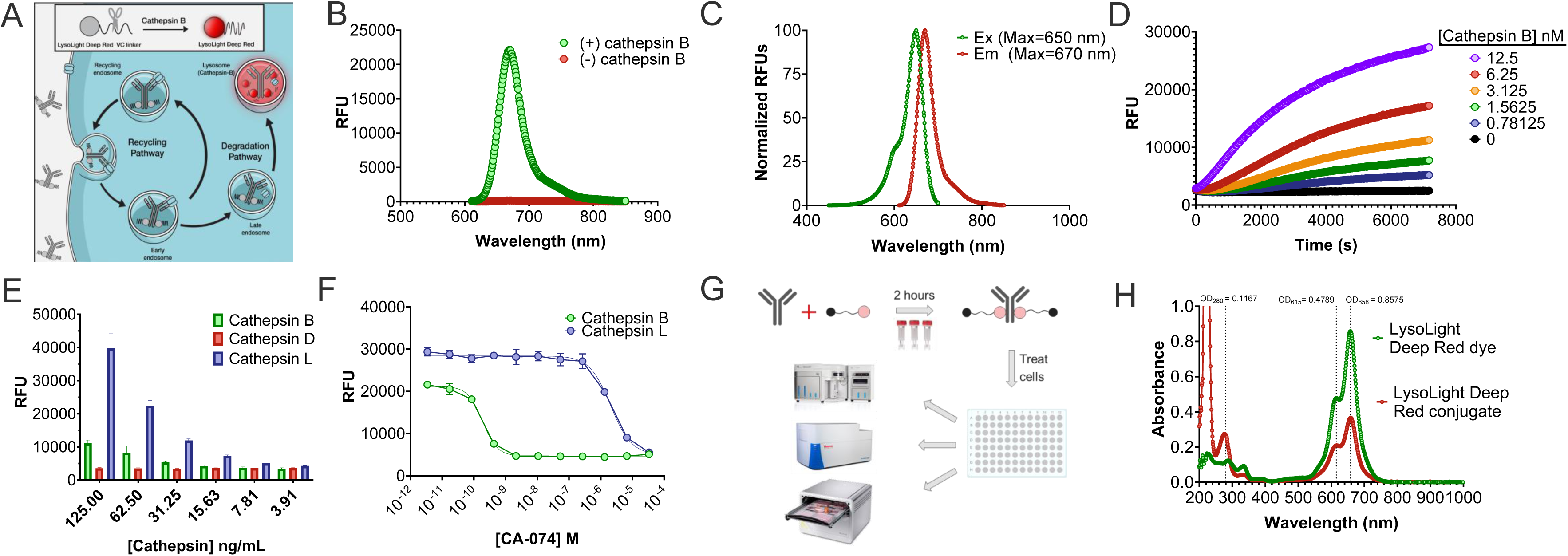
LysoLight Deep Red is a sensitive probe for Cathepsin B and L activity. A) Schematic describing internalization of LLDR antibody conjugates. B) Relative fluorescence intensity of LLDR in the presence (green) and absence (red) of recombinant Cathepsin B. C) Excitation (Ex – green) and emission (Em – red) profile of Cathepsin B cleaved LLDR. D) Relative fluorescence intensity of LLDR over time at various recombinant cathepsin B concentrations. E) Effect of Cathepsin B (green), D (Red), and L (blue) on LLDR. F) Inhibition of recombinant Cathepsin B and L with the Cathepsin B inhibitor, CA-074. G) Diagram describing the antibody conjugation workflow and applications. H) Absorbance spectra of LLDR (green) and LLDR antibody conjugate (red).

## Results

### LysoLight Deep Red is sensitive to cathepsin activity

LysoLight Deep Red (LLDR) was developed as a tool to monitor the catabolism of target proteins in the lysosome. Cathepsin B (CatB) is a lysosomal protease that plays a crucial role in this catabolic process, as its activity is regulated by the acidic environment of the lysosome[15–17]. In the field of antibody drug conjugates, researchers have utilized the selectivity of CatB by designing dipeptides with specificity for this protease. Utilizing a widely used ADC dipeptide, the valine-citrulline (VC) linker, we developed a probe that fluoresces only upon cleavage of the linker by cathepsin proteases.

We first studied the effectiveness of LLDR as a sensor for CatB activity in a cell-free assay using the LLDR carboxylate. In the absence of CatB, LLDR exhibited minimal fluorescence (Fig. 1B). However, upon the addition of recombinant CatB, a strong increase in fluorescence signal was observed (Fig. 1B). We characterized the spectral properties of LLDR after complete digestion by CatB and observed an excitation maximum at 650 nm and an emission maximum at 670 nm in 5 mM MES, pH 5.0 (Fig 1C). Additionally, we evaluated the sensitivity of LLDR to CatB activity and found that it could detect activity as low as 1.0 nM CatB after a 2-hour incubation (Fig 1D).

While the VC linker was initially believed to be specific for CatB, recent studies have shown that it can also be cleaved by other cathepsins [19,20]. To assess the promiscuity of LLDR, we compared the activity of CatB, CatL, and CatD for cleavage of LLDR. CatL exhibited significantly higher activity compared to CatB, while CatD showed little to no activity under the assay conditions (Fig 1E). Based on these findings and previous research, it appears that LLDR may not be specific for CatB alone but does show specificity for certain lysosomal cathepsins.

Next, we investigated the impact of inhibition on CatB and CatL activity using LLDR as a measure of enzymatic activity. We utilized the irreversible CatB inhibitor, CA-074, which has been shown to effectively inhibit CatB with IC50 values between 2 – 5 nM [21,22]. By titrating CatB and CatL with CA-074 for 1 hour followed by the addition of LLDR, we determined that the IC50 value for CatB inhibition by CA-074 was 0.188 nM, while the IC50 for CatL was nearly 10,000-fold higher at 2.0 μM (Fig 1F). This demonstrates the usefulness of CA-074 as an inhibitor for CatB activity and highlights its potential as a tool in investigating the specificity of LLDR for detecting lysosomal catabolism of target proteins in live cells.

### LysoLight™ Deep Red is a biologically active sensor to monitor lysosomal catabolism of a target protein

The cell-free experiments described above investigating the spectral properties and specificity were conducted using the free acid form of LLDR. To assess lysosomal degradation of the protein of interest, we functionalized LLDR with the amine reactive sulfodichlorophenol (SDP) ester to allow the conjugation of LLDR to any protein that possesses an accessible lysine residue, using a simple and robust conjugation method.

For example, optimized conditions for antibody labeling were to mix the dye and antibody in PBS at a 10-fold molar excess of dye to antibody at a final antibody concentration of 1 mg/mL under alkaline conditions. These conditions led to a degree of labeling (# of dye molecules per antibody molecule – DOL) between 1 and 2 after 2 hours of incubation. The DOL can easily be tailored depending on how long the reaction is allowed to proceed. Following incubation, the free excess dye is removed using a Zeba dye and biotin removal spin column equilibrated with PBS followed by addition and elution of the LLDR conjugate (Fig 1G). The absorbance spectrum of LLDR shows two peaks, one at 615 nm and the other at 650 nm. In PBS, the ratio of those two peaks changes between the free dye and the conjugate. However, dilution of either dye or conjugate at a ratio of 1:5 in Guanidine HCl lowers the peak intensity at 615 nm resulting in a similar ratio of the two peaks to allow for estimation of extinction coefficients and correction factors (Fig 1H).

To validate the utility of LLDR in live cell imaging-based assays, we conjugated LLDR to three well characterized therapeutic antibodies: trastuzumab (Her-2), cetuximab (EGFR), and rituximab (CD20)[23–25]. Since all three are of the human IgG1 subclass, LLDR was also conjugated to a human IgG1 isotype negative control. All antibodies had a degree of labeling between 1 and 2 with yields greater than 80%. We used the human breast cancer, Her2 positive, EGFR negative cell line, SKBR3 to assess internalization and lysosomal targeting of trastuzumab, cetuximab, and hIgG1 conjugates. A strong fluorescence signal was observed in SKBR3 cells with trastuzumab while minimal fluorescence in the two negative controls (cetuximab and hIgG1) (Fig 2A, top panel). Conversely, we used the EGFR positive/Her2 negative lung epithelial cell line, A431, to look at internalization and lysosome targeting of cetuximab conjugates. Here, little to no fluorescence was observed with the negative control conjugates, but positive signal was observed for cetuximab (Fig 2A, bottom panel). The signal intensity was quantified after subtraction of background observed with hIgG1 isotype control treatment (Fig 2B).

**Figure 2:**
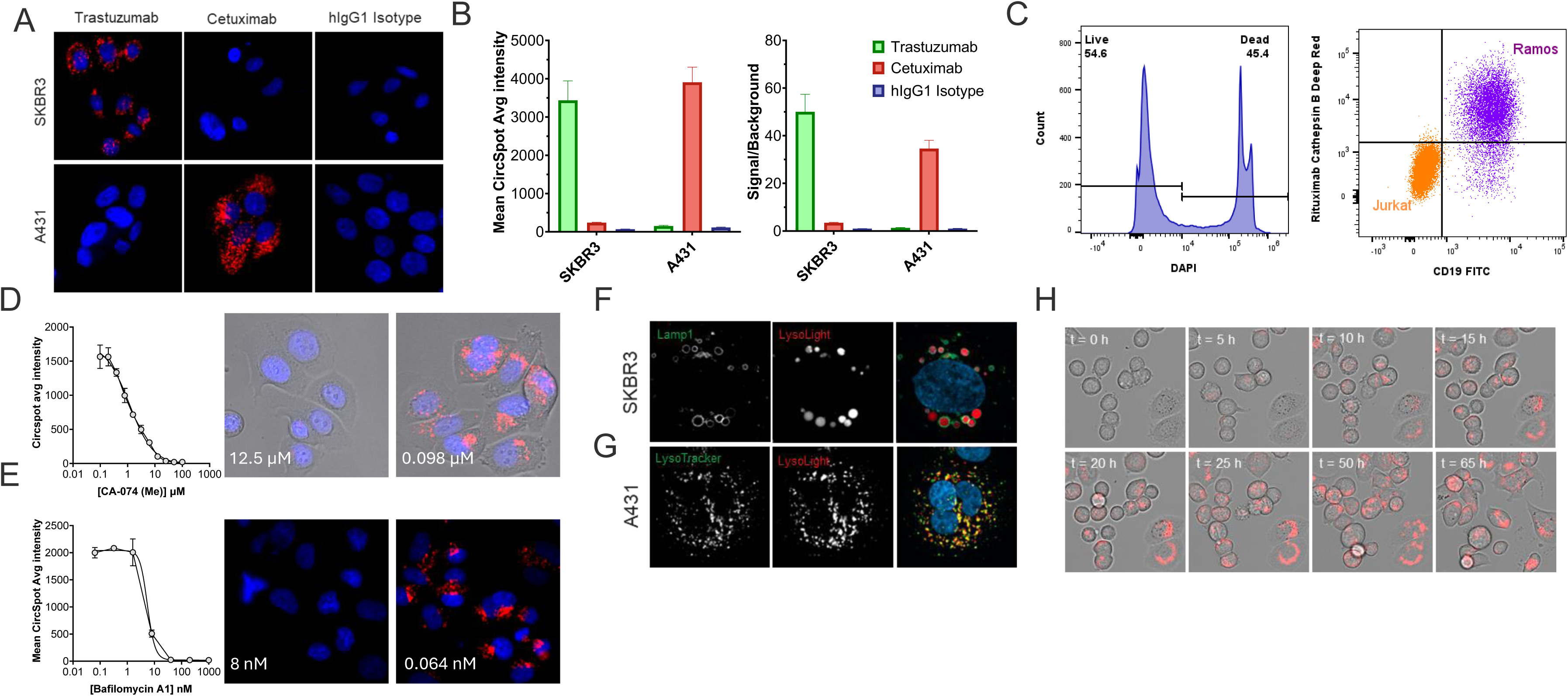
LysoLight Deep Red is a biologically active sensor to monitor lysosomal catabolism of a target protein. A) Treatment of SKBR3 or A431 cells with trastuzumab, cetuximab, or hIgG1 LLDR conjugates. B) Mean intensity of fluorescence signal from cells treated with LLDR conjugates. C) Flow cytometry measurements for Ramos (purple) and Jurkat (orange) cells treated with rituximab-LLDR. D,E) Mean average intensity of trastuzumab-LLDR treated SKBR3 cells titrated with CA-074 (Me) (D) or BafA (E) and representative images of high and low treatments. F) Images of SKBR3 cells treated with trastuzuamb-LLDR (Red) expressing Lamp-1 GFP (green). G) A431 cells treated with cetuximab-LLDR (Red) and LysoTracker Red (green). H) Still frames captured from 66 h time course of SKBR3 cells treated with trastuzumab-LLDR.

Next, we wanted to evaluate LLDR conjugates in flow cytometry applications using CD19 and CD20 positive Ramos cells and CD19/CD20 negative Jurkat cells. Cells were treated with rituximab-LLDR for 16 hours followed by staining with DAPI and CD19-FITC. CD19-positive Ramos cells strongly internalized and targeted rituximab-LLDR to the lysosome and while no lysosome targeting was observed in CD19-negative Jurkat cells (Fig 2C). Taken together, these results suggest that LLDR can be used to monitor lysosomal degradation of internalized therapeutic antibodies.

To assess selectivity of LLDR conjugates for Cathepsin B in cells, we used a methyl-ester form of CA-074 that is cell-permeable[26]. While CA-074 (Me) has a lower affinity for CatB, upon entering the cell, esterases activate the inhibitor by cleaving off the methyl ester. SKBR3 cells were treated with this inhibitor for 2 hours followed by addition of trastuzumab-LLDR. After 16 hours, images were captured and mean fluorescence intensity was determined. CA-074 (Me) completely abolished signal with an IC50 of 520 nM (Fig 2D) consistent with the LLDR primarily being a CatB substrate in cells.

If LLDR conjugates are being localized and degraded by the lysosome, then disruption of the lysosomal pH should disrupt the cathepsin activity and show a reduced response from LLDR. The vacuolar ATPase is the primary proton pump responsible for maintaining lysosomal pH and treatment with the vATPase inhibitor Bafilomycin A1 (BafA) results in lysosome alkalinization[27]. Treatment of SKBR3 cells with BafA, followed by treatment of trastuzumab-LLDR showed strong inhibition of LLDR signal (Fig 2E).

We next examined the subcellular localization of LLDR conjugates using common lysosomal markers. SKBR3 cells were transduced with CellLight Lysosomes-GFP, BacMam 2.0 baculovirus, to express GFP-Lamp1 fusion protein as a lysosome marker. Following transduction, cells were treated with trastuzumab-LLDR for 16 hours followed by confocal imaging. Nearly all lysosomes positive for Lamp1 were also positive for LLDR (Fig. 2F).

Furthermore, A431 cells treated with cetuximab-LLDR showed strong co-localization with LysoTracker Red, a cell-permeable fluorescent dye that stains acidic compartments within a cell (Fig 2G). Taken together, this data suggests that LLDR conjugates are specific for lysosomal localization and target degradation (Fig 2H).

### Lysolight Deep Red enables direct and sensitive assessment of LYTAC catabolism

As a lysosomal targeted degradation platform, monitoring lysosomal catabolism is of interest in LYTAC drug development. We therefore utilized the LLDR probe to examine the cellular fate of both the LYTAC and target. As a proof of concept, we generated LYTACs for targeted degradation of IgE. IgE targeting LYTACs are based on 3 components: 1) an IgE binding antibody (omalizumab), 2) an ASGPR binding ligand (modified GalNac) and a linker sufficient to allow co-engagement of IgE and ASGPR (Fig.3A)[4,28,29]. To enable direct comparison of ASGPR ligands, omalizumab-based LYTACs were generated with a fixed linker/ligand to protein ratio (LPR) of 2. For these proof-of-concept studies, ASGPR ligands with one (monovalent), two (bivalent) or three (trivalent) modified GalNac molecules per linker were selected to evaluate differential activity due to avidity-driven differences in LYTAC binding to the ASGPR trimer.

**Figure 3:**
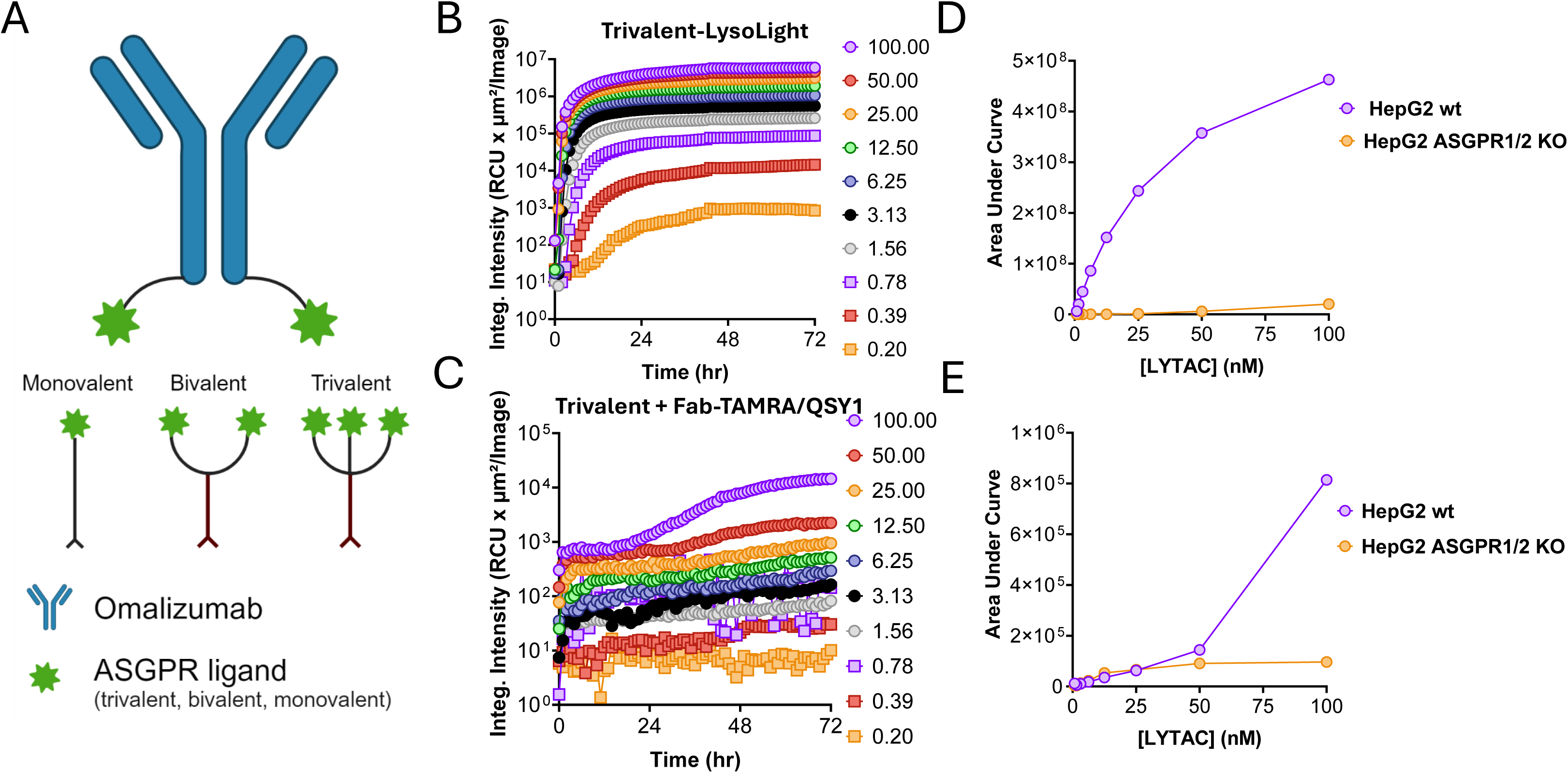
Monitoring LYTAC Catabolism Using Lysolight Conjugates. A) Schematic of LYTAC designs used in Figs 3-5. Omalizumab conjugated with monovalent, bivalent and trivalent ASGPR ligands or control conjugate without an ASGPR ligand (-ve). B) Integrated fluorescence intensity over time (72h) following incubation of LysoLight Deep Red labeled trivalent LYTAC at the indicated concentrations (nM) with wild type HepG2 cells. C) Experiment as described in (B) with LYTAC complexed with a quenched anti-IgG Fab. D,E) Area under the curve across the dose response for data shown in (B,C) and for the same experiment performed in ASGPR1/2 double knockout HepG2 cells.

LYTACs targeting the ASGPR internalizing receptor are internalized into an endosome where a decrease in pH and calcium levels results in dissociation of the LYTAC from the recycling ASGPR receptor into the endosome lumen and eventual delivery and catabolism of the LYTAC and target in the lysosome[2,4,30,31]. While pH-sensitive dyes exist to show internalization and late endosome/lysosome delivery of LYTACs and their targets, demonstrating catabolism of LYTAC and target has remained a challenge. One approach to assess catabolism of an internalized antibody utilizes an anti-human IgG F(ab)’2 reagent that is labeled with a fluorescent probe (TAMRA) and a quencher (QSY1)[32]. Proteolysis of the internalized antibody results in dequenching of fluorescence and therefore the fluorescent signal is a measurement of degradation of the internalized antibody. To assess the potential for each approach to measure LYTAC fate, we treated ASGPR expressing HepG2 cells or ASGPR1/2 knockout HepG2 cells with trivalent LYTAC either directly labeled with LLDR or with LYTAC precomplexed at 1:1 molar stoichiometry with anti-IgG F(ab’)2 conjugated with fluorophore and quencher (Fab-TAMRA/QSY1) across a dose response and monitored intracellular fluorescent signal by live cell imaging. LYTAC directly labeled with LLDR showed increased fluorescence over the 72 h time course across the full dose response from 100 nM to ∼200 pM (Fig 3B). In contrast, only the top two concentrations (50 and 100 nM) of the LYTAC + Fab-TAMRA/QSY1 showed a clear increase in signal across the time course (Fig 3C). By calculating the area under the curve for each time course for wild type and ASGPR1/2 knockout HepG2 cells, a signal to background ratio can be estimated for each detection reagent. The LYTAC-LLDR treatment showed signal to background ratios (S/B) of 59 and 22 at 50 nM and 100 nM doses (Fig. 3D) versus a S/B of only 1.5 and 8.5 for the LYTAC + Fab-TAMRA/QSY treatment at the same concentrations (Fig. 3E). These results indicate that LLDR is a suitable probe to directly measure delivery and catabolism of LYTAC in the lysosome with superior S/B to an alternative measurement of catabolism.

### ASGPR ligand valency influences IgE degradation activity of omalizumab-based LYTACs

Having demonstrated the utility of LLDR to measure LYTAC catabolism in the lysosome, we next sought to determine if the probe is suitable for measuring target catabolism using LYTACs that are expected to vary in activity due to the number of ASGPR ligands. As a minimal test set, we compared the activity of an ASGPR ligand valency series consisting of monovalent (1 ASGPR ligand per linker at LPR of 2 = 2 total ASGPR ligands per LYTAC), bivalent (4 ASGPR ligands per LYTAC) and trivalent (6 ASGPR ligands per LYTAC) omalizumab-based LYTACs (see Fig 3A). To compare activity across the valency series, IgE conjugated with LLDR was mixed at 1:1 stoichiometry with LYTAC across a dose response ranging from 100 nM to 0.14 nM and fluorescence was measured by live cell imaging over 72 h. As expected, the trivalent ligand showed the highest activity followed by bivalent and monovalent ligands (Fig. 4A-C).

**Figure 4:**
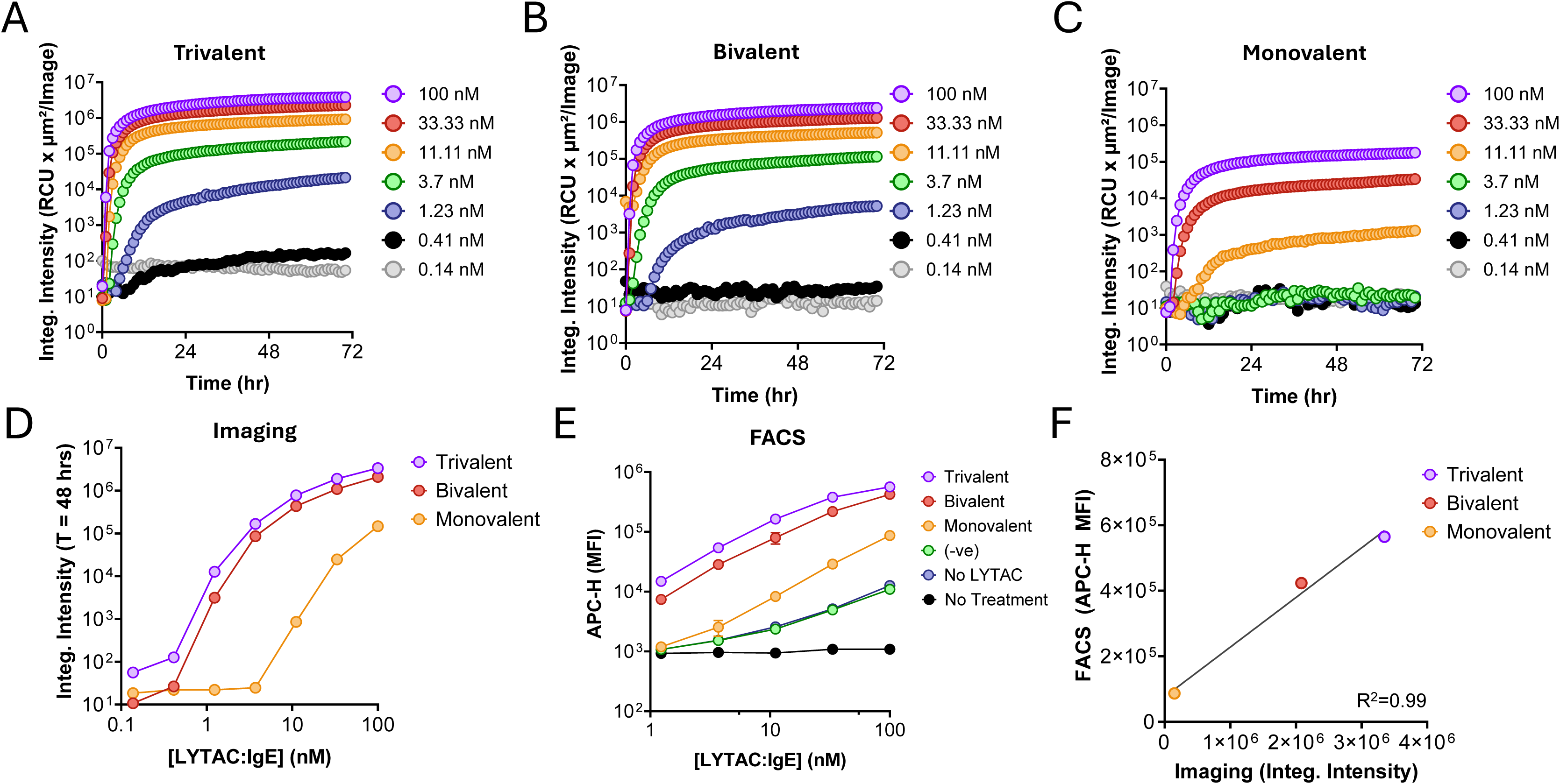
Assessing IgE Degradation Activity by omalizumab-based LYTACs. A-C) Integrated fluorescence intensity over time (72h) following incubation of IgE-LysoLight Deep Red and LYTAC at the indicated concentrations (nM) with HepG2 cells. D) Integrated fluorescence intensity at 48 h across the dose response shown in A-C. E) Fluorescence measured by flow cytometry (FACS) across IgE-LysoLight Deep Red dose response with the indicated treatments in HepG2 cells. F) Linear correlation between imaging (D) or FACS (E) measurements for indicated treatments at 100 nM.

The trivalent and bivalent ligands showed signal above background down to 1.23 nM (Fig. 4A,B). The less active monovalent ligand showed sensitivity to 11.1 nM (Fig. 4C). Across the dose response, the trivalent ligand performed slightly better than the bivalent ligand while the monovalent ligand showed diminished potency of >10-fold (Fig. 4D). To confirm the results from live cell imaging, we performed a similar dose response (100-1.23 nM) and analyzed by flow cytometry 24 h after treatment. Consistent with the imaging results, the trivalent ligand showed slightly more activity than the bivalent ligand which was much more active than the monovalent ligand (Fig. 4E). To confirm that activity was due to ASGPR/ligand engagement and not a non-specific property of omalizumab bound to IgE-LLDR, we generated omalizumab reacted with maleimide without a linker-ligand (-ve). We observed no activity above that of the no LYTAC (IgE-LLDR) control (Fig.4E). Finally, we observed a strong correlation between measurements of LYTAC activity using the imaging and flow cytometry approaches demonstrating the reproducibility of activity estimates using orthogonal measurements (Fig. 4F).

### Assessment of IgE degradation activity by omalizumab-based LYTACs in primary human hepatocytes

LYTACs utilizing ASGPR ligands are expected to be primarily directed to hepatocytes *in vivo* as has been observed in the siRNA delivery field[29]. While the HepG2 hepatocellular carcinoma cell line expresses ASGPR, it was possible that the fate of internalized IgE might be different between HepG2 and human hepatocytes leading to differential catabolic activity. In addition, the ASGPR level is reported to be >10X lower in HepG2 cells compared to hepatocytes and hepatocytes express detoxifying enzymes that are not present in HepG2[33–35]. We therefore sought to determine if LYTAC-mediated IgE-LLDR catabolism in the lysosome can be assessed in primary hepatocytes by both live cell imaging and flow cytometry.

Internalization and lysosome catabolism of IgE-LLDR was stimulated by treatment with LYTACs (100 nM) displaying trivalent, bivalent and monovalent ASGPR ligands but not with omalizumab (-ve) in primary human hepatocytes over a 24 h time course as measured by live cell imaging (Fig.5A). An endpoint of 24 h was chosen for primary hepatocytes, as opposed to 72 h for HepG2 cells, due to treatment-independent decreased hepatocyte cell viability observed at later timepoints. As observed in HepG2 cells, LYTACs with a trivalent ASGPR ligand display were most active followed by bivalent and monovalent ligands (Fig.5B). However, primary hepatocytes showed less differentiation between the bivalent ligand and monovalent ligands compared to the same treatments in HepG2 cells. To confirm that primary hepatocytes show less differentiation between ligands than HepG2 cells, a flow cytometry-based assay was run under the same treatment conditions as performed in HepG2 cells (Fig. 5C). As for HepG2 cells, strong concordance was observed between the imaging-based and flow cytometry-based measurements for activity in primary hepatocytes (Fig. 5D). To allow for comparison between cell types, activity for the bivalent and monovalent ligands as measured by flow cytometry were normalized to the trivalent activity. The bivalent and monovalent ligands showed 82.9% and 54.8% activity (Fig. 5C) relative to the trivalent ligand in primary hepatocytes vs. 75.0% and 15.4% in HepG2 cells (Fig. 4E). These results indicate that primary hepatocytes are less sensitive to changes in valency than the HepG2 cell line and therefore that lower valency ligands might show higher than expected *in vivo* activity than predicted by assessment in HepG2 cells alone. Taken together, we conclude that the LLDR probe allows for sensitive and quantitative assessment of LYTAC catabolic activity in both cell lines and primary cells by multiple approaches.

**Figure 5:**
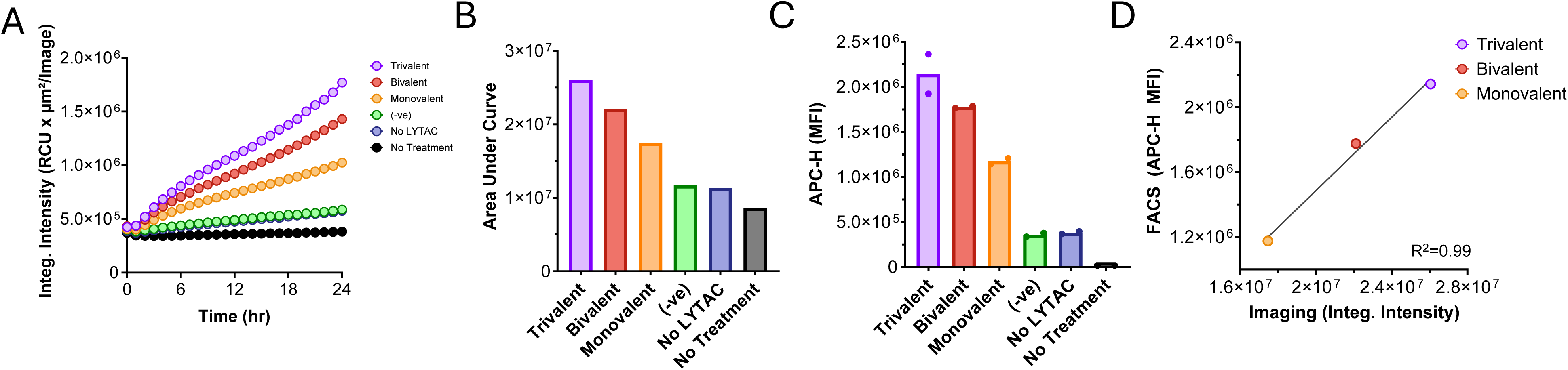
Assessing IgE Degradation Activity by omalizumab-based LYTACs. A,B) Integrated fluorescence intensity over time (24h) (A) and areas under the curves (B) following incubation of IgE-LysoLight Deep Red and LYTAC at 100 nM each with primary human hepatocytes. C) Same experiment as in A,B measured by flow cytometry at 24h. D) Linear correlation between imaging (C) or flow cytometry (D) measurements for indicated treatments at endpoint (24h).

## Discussion

This study demonstrates the application of LLDR as a sensitive tool for monitoring cathepsin activity and lysosomal catabolism of target proteins. LLDR showed minimal fluorescence in the absence of Cathepsin B (CatB), but a strong increase in fluorescence upon the addition of recombinant CatB, confirming its utility as a sensor for lysosomal proteases. The spectral properties of LLDR and its ability to detect CatB activity at concentrations as low as 1.0 nM further underscore its sensitivity.

While LLDR was initially designed with the valine-citruline (VC) linker for CatB specificity, our findings reveal that LLDR also responds to other cathepsins, such as CatL with varying efficiencies. The inhibition studies using the CatB-specific inhibitor CA-074 validated LLDR specificity for CatB, with an IC50 value of 0.188 nM, highlighting its potential for investigating lysosomal catabolism in live cells.

Functionalization of LLDR with the amine-reactive sulfodichlorophenol (SDP) ester enabled its conjugation to proteins with accessible lysine residues, allowing for the assessment of target protein degradation. Optimized conjugation conditions yielded LLDR conjugates with a degree of labeling (DOL) between 1 and 2, with high yields. Validation in live cell imaging assays showed strong fluorescence signals with therapeutic antibodies trastuzumab, cetuximab, and rituximab in target-positive cell lines, confirming the utility of LLDR to report lysosomal degradation of target proteins. Flow cytometry applications further supported these findings, showing specific internalization of rituximab-LLDR in CD19 and CD20 positive Ramos cells.

The application of LLDR in monitoring lysosomal catabolism in LYTAC drug development was also demonstrated. LLDR enabled sensitive and quantitative assessment of LYTAC catabolic activity in ASGPR-expressing HepG2 cells and primary human hepatocytes. The superior signal-to-background ratios observed with LLDR compared to another catabolism measurement that is specific only for human antibodies (a quenched anti-human IgG F(ab’)2) suggests that LLDR can be applied to monitor catabolism of a broader range of proteins of interest than existing approaches. Additionally, LLDR was useful in evaluating the impact of ASGPR ligand valency on LYTAC activity, showing that higher valency ligands exhibited greater activity, with strong correlation between imaging and flow cytometry measurements.

In primary human hepatocytes, LLDR successfully measured LYTAC-mediated IgE catabolism, demonstrating its applicability in both cell lines and primary cells. The results indicated that primary hepatocytes are less sensitive to changes in ligand valency compared to HepG2 cells, suggesting that lower valency ligands might show higher than expected *in vivo* activity.

In conclusion, LLDR is a robust and versatile tool for monitoring cathepsin activity and lysosomal catabolism of target proteins. LLDR is a highly sensitive, specific, and broadly applicable probe to measure lysosomal degradation pathways with application in therapeutic lysosomal-targeted delivery/degradation platforms, such as conventional ADCs and LYTACs. The performance in multiple cell types and measurement approaches further establishes LLDR as a critical tool for both biological and therapeutic research applications.

## Methods

### Cell free LysoLight Deep Red Assay

Recombinant Cathepsin B was diluted to 10 µg/mL in 25 mM MES, 5 mM DTT, pH 5.0, incubated at room temperature for 15 minutes and then further diluted to final 2x concentration in 25 mM MES, pH 5.0. 20 µL of 5 µM LysoLight deep red was added in triplicate to wells of a black 384 well plate. 15 µL Cathepsin B or buffer was then added to 15 µL LysoLight and allowed to react for 2 hours at room temperature. To determine emission maximum, an emission scan was performed holding the excitation constant at 580 nm, while to determine the excitation maximum, an excitation scan was performed holding the emission constant at 740 nm. All other cell free assays were performed using Tecan Spark’s kinetic mode taking reading every 1 minute for 2 hours. In the case of inhibition studies, 15 µL of inhibitor was added to 15 µL Cathepsin B for 1 hour prior to addition of 15 µL LLDR.

### LysoLight Deep Red conjugation and purification

Antibodies were prepared for labeling at a concentration of 2.5 mg/mL in phosphate buffered saline (PBS). 50 µL of 1 M Sodium Bicarbonate (pH 8.4) was subsequently added to 400 µL of prepared antibody to raise the pH. Finally, 50 µL of LLDR (diluted to 2 mg/mL) was added to the antibody cocktail and reacted at 25°C for 2 hours with shaking on an orbital shaker at 550 RPM in the dark. Antibodies were purified with 2 mL Zeba™ dye and biotin removal columns equilibrated with PBS. Briefly, columns were spun at 1,000 g for 2 minutes to remove storage buffer. Columns were then equilibrated with PBS exchange buffer and spun for an additional 2 minutes. Antibody conjugates were added to the center of the column and spun at 1,000 g for 2 minutes to elute purified material. Finally, purified conjugates were spun at 14,000 g for 10 minutes to remove any aggregates or precipitation and stored at 4°C until use. Following purification, conjugates were quantified by diluting 1:4 in 8 M guanidine HCl and measuring the absorbance at 280 nm and 658 nm. The protein concentration in M and the degree of labeling (DOL) were calculated using the following formulas, where ε is the extinction coefficient of the protein (∼210,000 cm^-1^M^-1^), ε’ is the extinction coefficient of LLDR in 8 M Guanidine HCl (344,500 cm^-1^M^-1^), and CF is the correction factor for LLDR at 280 nM (0.187).

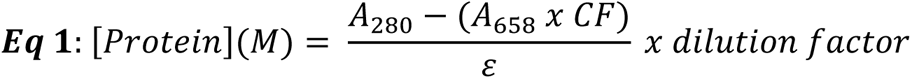

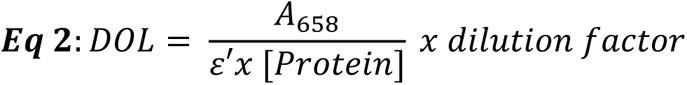

### Treatment of cells with LysoLight Deep Red conjugates

SKBR3 or A431 cells were seeded at a density of 7,500/well in complete media supplemented with 10% FBS and allowed to recover for 16 h. Media was removed and replaced with fresh media. 10x stocks of LLDR conjugates were prepared by diluting to 10 μg/mL in Live Cell Imaging Solution (LCIS). 10 µL of 10x stocks were added to cells in triplicate and grown at 37°C, 5% CO_2_, 80% relative humidity for 16 hours. Following conjugate treatment, Hoechst 33342 was added at a dilution of 1:8,000 for 15 minutes stored under the conditions above. Cells were then imaged on the Thermo Scientific™ CellInsight™ CX7 LZR Pro. 200 cells/well were segmented on the nucleus and then mean intensity of each cell was averaged over all 200 cells. Exposure settings were matched across all positive and negative controls. For time course studies, cells were treated as above, and then imaged on the EVOS M7000 with OSI2 to control temperature, CO_2_/O_2_, and humidity, for 65 hours taking images every 15 minutes.

For live cell inhibition studies, drugs were diluted as 10x stocks in LCIS followed by 2-fold serial dilutions. Cells were prepared as above followed by 10 μL addition of the drug along with a no inhibitor control in triplicate. Cells were incubated under standard tissue culture conditions for 2 hours followed by the addition of trastuzumab-LLDR at a final concentration of 1 μg/mL.

For co-localization experiments with LysoTracker Red, cells were treated as above. Following 16 h incubation with antibody conjugates, cells were treated with 50 nM LysoTracker Red in complete media for 30 minutes. Images were captured on the EVOS M7000 using the Texas Red (LysoTracker Red) and Cy5 (LysoLight Deep Red) filter sets.

### LYTAC Generation

Omalizumab containing an engineered cysteine substitution was expressed in Expi293 cells (Thermo Fisher Scientific) and purified using a Protein A mAb Select SuRe column, followed by cation exchange chromatography on an SP-HP column with an AKTA Go Protein Purification System (Cytiva Life Sciences). The purified protein was characterized using intact mass analysis for mass confirmation, SEC-HPLC (>95% main peak), and SDS-PAGE for purity. Omalizumab was conjugated to the maleimide-functionalized linker-ligands via site-specific conjugation of the engineered cysteine. Briefly, omalizumab was buffer exchanged into 100 mM HEPES, 50 mM NaCl, 1 mM EDTA, pH 7.2 (HBSE buffer), and reduced with 20 molar equivalents of 0.5 M TCEP (Bond-Breaker™ TCEP Solution, Thermo Fisher Scientific) at 37 °C for 1 h. The reducing agent was then removed using Zeba™ spin desalting columns, equilibrated with HBSE, and the reduced antibody was re-oxidized with 40 molar equivalents of dehydroascorbic acid at room temperature for 2 h. The re-oxidized omalizumab was then incubated with 5 molar equivalents of each linker-ligand (monovalent, bivalent or trivalent) at room temperature for 1-2 h. The reactions were purified by size exclusion chromatography in phosphate buffered saline pH 7.5.

The identity and degree of conjugation of the purified conjugates were determined via liquid chromatography-mass spectrometry (LC-MS). All conjugates were >95% monomeric as determined by size exclusion high performance liquid chromatography (SEC-HPLC).

### LYTAC catabolism evaluated by LysoLight Deep Red or anti-IgG F(ab’)2-TAMRA-QSY1

Wild-type (WT) and ASGPR1/ASGPR2 double knockout (ASGPR1/2 KO) HepG2 cells were cultured in EMEM (ATCC) supplemented with 10% FBS and 1X Pen/strep and maintained at 37°C at 5% CO_2_. The ASGPR1/2 KO cells were generated by CRISPR/Cas9 and subsequent clonal expansion. HepG2 cells were seeded in 96-well plates 48h prior to treatment with LYTAC molecules, at a density of 50,000 cells/well. On the day of the experiment, media was removed from cells and 45 µl of EMEM+1/100 Human TruStain FcX blocking solution (Fc Block, BioLegend) was added. While cells were incubating with Fc block, LYTAC directly conjugated to LysoLight DeepRed (100 nM) (LYTAC-LLDR) or LYTAC precomplexed to Fab(2)-TAMRA/QSY (100 nM each) (LYTAC+Fab-T/Q) were serially diluted (2-fold) in a 96 well plate. After cells incubated in Fc Block for 10 minutes, 45 μl was transferred from the serial dilution plate to cells. Plates were incubated for 72h at 37°C. After incubation, LYTAC uptake was measured by flow cytometry (referred to as FACS) or Incucyte live-cell imaging.

For FACS evaluation, cells were rinsed twice with Dulbecco’s phosphate buffered saline (DPBS, Gibco), lifted with 0.25% Trypsin/EDTA (Gibco), transferred to conical bottom 96-well plates and resuspended in 100 μL of FACS Buffer with BSA (Rockland Immunochemicals). Mean fluorescence intensity in the APC channel was measured using a Novocyte Advanteon flow cytometer.

For live cell imaging experiments, LysoLight Deep Red fluorescence was measured in the red channel every hour using Incucyte Live Cell Analysis System (Sartorius) over the course of the 72 h incubation.

### HepG2 In Vitro IgE-LysoLight Catabolism Assay

HepG2 cells (WT or ASGPR1/2 knockout) were cultured in EMEM (ATCC) supplemented with 10% FBS and 1X Pen/strep and maintained at 37°C and 5% CO_2_. Cells were seeded in 96-well plates 48h prior to LYTAC treatment at a density of 50,000 cells/well. On the day of the experiment, 200 nM of LYTAC molecules were precomplexed with equimolar IgE conjugated to LysoLight Deep Red (IgE-LLDR) for 30 minutes in complete media. During the precomplexing step, cells were removed from the incubator and media was replaced with 45 µl of EMEM+1/100 Fc block. While cells were incubating with Fc Block (10 minutes), the precomplexed LYTAC:IgE-LLDR mix was serially diluted in 96-well plate (3-fold), after which 45 μl of diluted stocks were added from serial dilution plate to cells with media+Fc blocking solution.

Cells were incubated with LYTAC:IgE-LLDR mixture for 3h at 37°C and analyzed by FACS as described above. Alternatively, cells were incubated with LYTAC:IgE-LLDR for 72h and analyzed by live-cell imaging as described above.

### Human Primary Hepatocyte In Vitro IgE-LysoLight Catabolism Assay

24-well plates were coated with 500 µl of Collagen I per well (ThermoFisher, A1048301) and incubated for 3h at room temperature. After 3h, wells were rinsed twice with DPBS and dried overnight. Human primary hepatocytes cells (Xenotech, HPCH05+) were thawed and the collagen coated plates were seeded at a density of 1.2×10^6^ cells/well, according to manufacturer’s instructions. The cells were incubated at 37°C and 5% CO_2_ for 2h to allow attachment. The OptiThaw media was aspirated, and OptiCulture media was added to the wells and incubated 16h. 100 nM of LYTAC was precomplexed with 100 nM IgE-LLDR in OptiCulture media for 30 minutes in a separate 24-well plate. During incubation of the precomplex, media was aspirated from cells and 165 µl of EMEM+1/100 Fc Block was added and incubated for 10 minutes. 165µl of precomplexed LYTAC:IgE-LLDR was transferred to the plate containing cells and Fc Block and incubated for 24h at 37°C. LysoLight Deep Red fluorescence was measured by Incucyte live cell imaging by imaging in the red channel every hour for 24 h. After 24 h incubation, cells were lifted and analyzed by flow cytometry as described above.

---

**Fig. 4; Supplemental 1:**
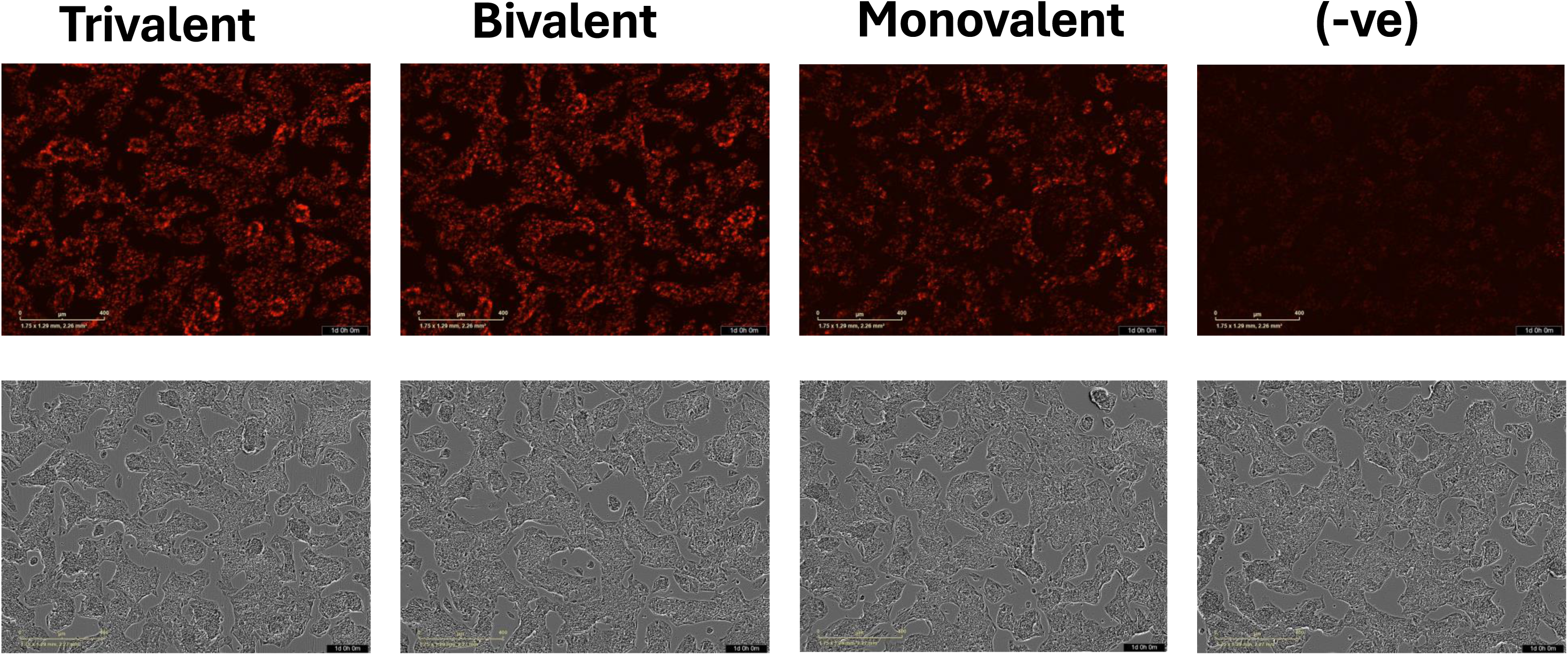
Representative images from figure 4 A-C from 24 hrs post-treatment with the indicated LYTAC and IgE-LLDR. Images were acquired on an Incucyte S3 Live-Cell Analysis Instrument at 100X total magnification.

## Notes

### Competing Interest Statement

RWH, YZH AC and CL are employees of ThermoFisher Scientific. VBV, CK, NP, TC, DL, SR, KHD, ET and MJS are employees of Lycia Therapeutics.

